# Testing the moderation of quantitative gene by environment interactions in unrelated individuals

**DOI:** 10.1101/191080

**Authors:** Rasool Tahmasbi, Luke M. Evans, Eric Turkheimer, Matthew C. Keller

## Abstract

The environment can moderate the effect of genes – a phenomenon called gene-environment (GxE) interaction. There are two broad types of GxE modeled in human behavior – qualitative GxE, where the effects of individual genetic variants differ depending on some environmental moderator, and quantitative GxE, where the additive genetic variance changes as a function of an environmental moderator. Tests of both qualitative and quantitative GxE have traditionally relied on comparing the covariances between twins and close relatives, but recently there has been interest in testing such models on unrelated individuals measured on genomewide data. However, to date, there has been no ability to test quantitative GxE effects in unrelated individuals using genomewide data because standard software cannot solve nonlinear constraints. Here, we introduce a maximum likelihood approach with parallel constrained optimization to fit such models. We use simulation to estimate the accuracy, power, and type I error rates of our method and to gauge its computational performance, and then apply this method to IQ data measured on 40,172 individuals with whole-genome SNP data from the UK Biobank. We found that the additive genetic variation of IQ tagged by SNPs increases as socioeconomic status (SES) decreases, opposite the direction found by several twin studies conducted in the U.S. on adolescents, but consistent with several studies from Europe and Australia on adults.

---

The effects of genes do not exist in a vacuum; they are likely to be influenced by the environmental background to various degrees. Understanding such GxE interactions has been a major focus of disease and behavioral genetic research over the past twenty years. Much of this research has investigated qualitative G×E effects using a candidate gene approach, such that the effects of specific genetic polymorphisms chosen a-priori based on biological hypotheses were modeled as a function of environmental moderators (e.g., [1]). However, concerns of high false positive rates [2], a history of poor replication [3], and the realization that individual genetic effect sizes are typically very small [4] has cast doubt on the utility of candidate gene-by-environment interaction studies. An alternative approach is to ask whether genetic effects across the genome change, on average, across an environmental moderator [5]. Qualitative GxE effects (see Supplemental Text) manifest as a non-unity genetic correlation between the same trait at different levels of an environment. Tests of such qualitative G×E effects have long been employed in samples of close relatives and twins [6], but have recently been tested among unrelated individuals using genome-wide SNP data, instantiated in the popular GCTA software using a mixed linear effects approach [7].

Maximum likelihood methods using close relatives have also been used to test quantitative GxE effects, in which genetic or environmental variance components change across the level of moderator [8]. Twin analyses of depression [1] [9] [10] [11] [12] [13], schizophrenia and bipolar disorder [14], alcohol and drug use and abuse [15] [16] [17], and others traits [18] [16] [17] have shown that the genetic and/or environmental variation underlying human behavior is often non-constant across different environments. Perhaps the best known example of this approach was Turkheimer's [19], finding that the additive genetic variance, V_A_, of IQ was lower for low SES than high SES individuals, which had also been reported previously [20] [21] [22] [23] [24] [25] [26]. This study prompted multiple follow-up studies, with some replicating the original finding and others not (Table S1).

Testing for quantitative GxE effects in unrelated individuals is important because close family members share environmental and non-additive genetic factors that, in combination, can lead to serious biases in estimates of additive genetic variation [27] [28] [29]. Furthermore, much more genome-wide data is available to researchers than twin/family data, and this is especially so for rare disorders. To date, however, there has been no ability to directly test quantitative GxE effects in a unified modeling approach using genome-wide SNPs in unrelated samples. Instead, to investigate changes in the genetic variance tagged by SNPs across a moderator, samples have been binned at different levels of the moderator, with genetic variance or SNP-heritability assessed separately in each group [30] [31]. Unfortunately, such an approach loses power compared to an approach that models all the data simultaneously, assumes that variances do not change as a function of the moderator within bins, and it make it difficult to test functionally different forms of possible interactions. Furthermore, if heritability (rather than additive genetic variance) is estimated separately per bin, it is implicitly assumed that variances are equal across bins, whereas what is often of interest is whether the absolute magnitude of genetic or environmental variation changes. In this paper, we introduce a model for testing quantitative GxE effects in unrelated samples using genome-wide SNPs and assess its accuracy by simulation. We then apply this method to a sample of 40,172 individuals in the UK Biobank to understand whether and how genetic variation underlying IQ changes as a function of SES in this population.

To model unrelated individuals, let

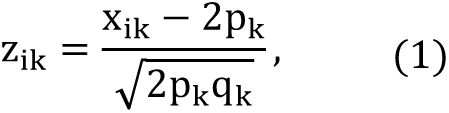

be the standardized genotype for individual i and variant k, where x_ik_ ∈ {0,1,2} and p_k_ is the allele frequency for the kth variant. Denote by V_P_ the phenotypic variance component and V_E_ non-genetic variance component, such that V_P_ = V_A_ + V_E_.

We write the quantitative gene by environment interaction model for individual i as

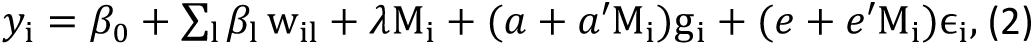

where *β*_1_’s are coefficients corresponding to w_il_ covariates, M_i_ is the standardized moderator and *λ* its effect, 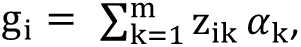 with the coefficients *α*_k_’s representing genetic effects for each of m SNPs assumed to be drawn from a normal distribution with mean zero and variance 1/m, and ε_i_’s representing error effects drawn from a standard normal distribution and independent from *α*_k_. The *a* and *e* coefficients represent the importance of additive genetic and environmental factors, respectively, while the *a*′ and *e*′ coefficients represent the degree to which the additive genetic and environmental influences change as a function of the moderator, *M*. In this (full) model, denoted Model 1, the additive and error variances are V_A_ = (*a* + *a*′M)^2^ and V_E_ = (*e* + *e*′M)^2^, which change as a function of moderator. Purcell [8] used a similar model for twin data.

From within this framework, we can define other models where V_A_ is constant but V_E_ changes as a function of moderator by setting *a*′ = 0 (Model 2), where V_A_ changes as a function of moderator but V_E_ is constant by setting *e*′ = 0 (Model 3), and where both V_A_ and V_E_ are constant by setting *a*′ = *e*′ = 0 (Model 4). Model 4 is the same as the base REML model instantiated in GCTA, such that V_A_, V_E_ and *h*^2^ are constant, but is useful for comparison and hypothesis testing. We also define a final model, Model 5, where V_A_ and V_E_ change as a function of moderator, but *h*^2^ is constant, by constraining *e*′ = *ea*′/*a* (note that setting *e*′ = *a*′ does not accomplish this; see Supplement). This model is useful for testing if changes in V_A_ and V_E_ are more parsimoniously explained by a change in V_*p*_. To the best of our knowledge, Model 5 or equivalent models, where the proportionate changes of V_A_ and V_E_ are constrained to be equal, have not been developed or tested in models designed for twin/family data (e.g., [8]). The best model for fitting the data can be determined based on formal hypothesis tests or, for models that are not nested, on the AIC/BIC fit criteria.

For Model 1,

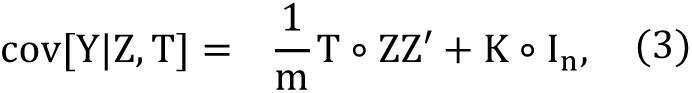

where *Y* = [*y*_1_,…,*y*_*n*_]′ is the column vector of phenotypes and *l*_*n*_ is the identity matrix of size *n*, *Z* = [*z*_*ik*_] is the standardized genotype matrix, *T* = [(*a* + *a*′*M*_*i*_)(*a* + *a*′*M*_*j*_)], *K* = [(*e* + *e*′*M*_*i*_)(*e* + *e*′*M*_*j*_)], and the operator ° is Schur product (or matrix element-wise product). (Details for computing the covariance matrices for all five models are shown in the Supplement). We assume Y = [y_1_,…, y_n_]′ follows a normal distribution with mean β_0_ + Σ_1_β_1_ w_il_ + λM_i_ and the covariance matrix estimated for each model. If we let 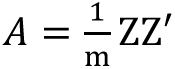 be the estimated genetic relationship matrix (GRM) from whole genome SNP data, equation (3) becomes

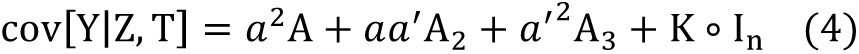

where *A*_2_ and *A*_3_ are additional GRMs that are functions of both the moderator and A (see Supplement for details). As can be seen, the coefficient of the second term (*aa*′) is a function of the first and third term, which makes equation (3) a constrained covariance matrix. REML (implemented by GCTA) can deal with multiple GRMs if their coefficients are independent of each other, but that is not the case here. If these three GRMs (A, A_2_, A_3_) are entered into GCTA, it will estimate a coefficient of the second term that is not constrained to equal *aa*′. Here, we maximize the log-likelihood function using parallel constrained optimization (see the definition of matrix *V* in [32], page 77) assuming that the phenotypes follow a multivariate normal distribution (see Methods).

We ran a comprehensive set of simulations (Method section) to investigate the performance of the proposed method. Phenotypes were simulated from each of the 5 models with 6 different sets of parameters (Table S2) and different sample sizes. The results are shown in Tables S3-S6. The biases of the estimated parameters were not statistically significant from zero. The simulation results for type I error are presented in Supplementary Figures S1-S9 and Tables S7-S12, and show no inflation of type-I error rates. Figure 1 presents the statistical power for testing *a*′ = 0 in Model 3, and shows 80% power for detecting a 5% increase in V_A_ for every standard deviation increase in the moderator (when a=.63 and a’=.04) once sample sizes are above 8000. The power for a given parameter differs across models (see Supplementary Tables S7-S12 and Supplementary Figures S10-S17), and is lower in models attempting to estimate more parameters due to correlations between the estimates. For example, 80% power for detecting *a*′ in Model 3 requires a sample size of 1000, but due to the correlation between estimates of *a*′ and *e*′, requires a sample of size 6000 in Model 1.

**Figure 1.**
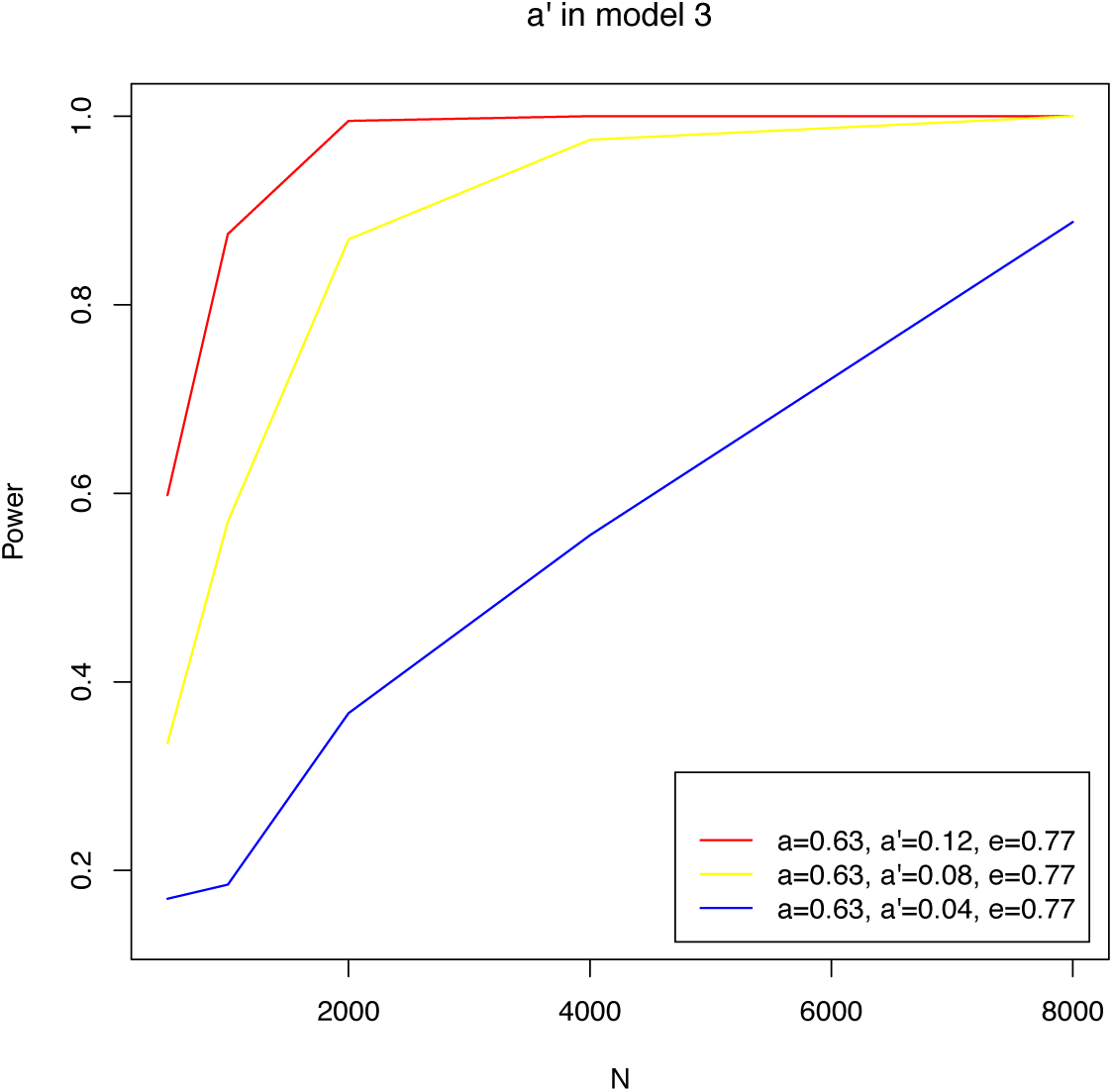
Power plot for testing *a*′ = 0 in model 3 with different set of parameters.

Figure 2 shows results from a sensitivity analysis, where the data are simulated from Model 1 and the parameters are estimated from Models 1-5, to show the effects of model misspecification on parameter estimates. Estimates are unbiased when the correct model is used, but *a*′ is overestimated and *a* underestimated when *e*′ is incorrectly dropped, and *e*′ is overestimated and *e* underestimated when *a*′ is incorrectly dropped. When both *a*′ and *e*′ are incorrectly dropped, such as would occur using the traditional approach and not allowing for moderation of V_A_ or V_E_, estimates for *a* are unbiased but estimates for *e* are overestimated, leading to underestimation of *h*^2^. Figures S18-S20 show similar results where data are simulated from models 2, 3, 4, and 5 respectively (see also Tables S18-S21). Overall, our results indicate that estimates are unbiased when the correct model is chosen but can be biased to various degrees under model misspecification.

**Figure 2.**
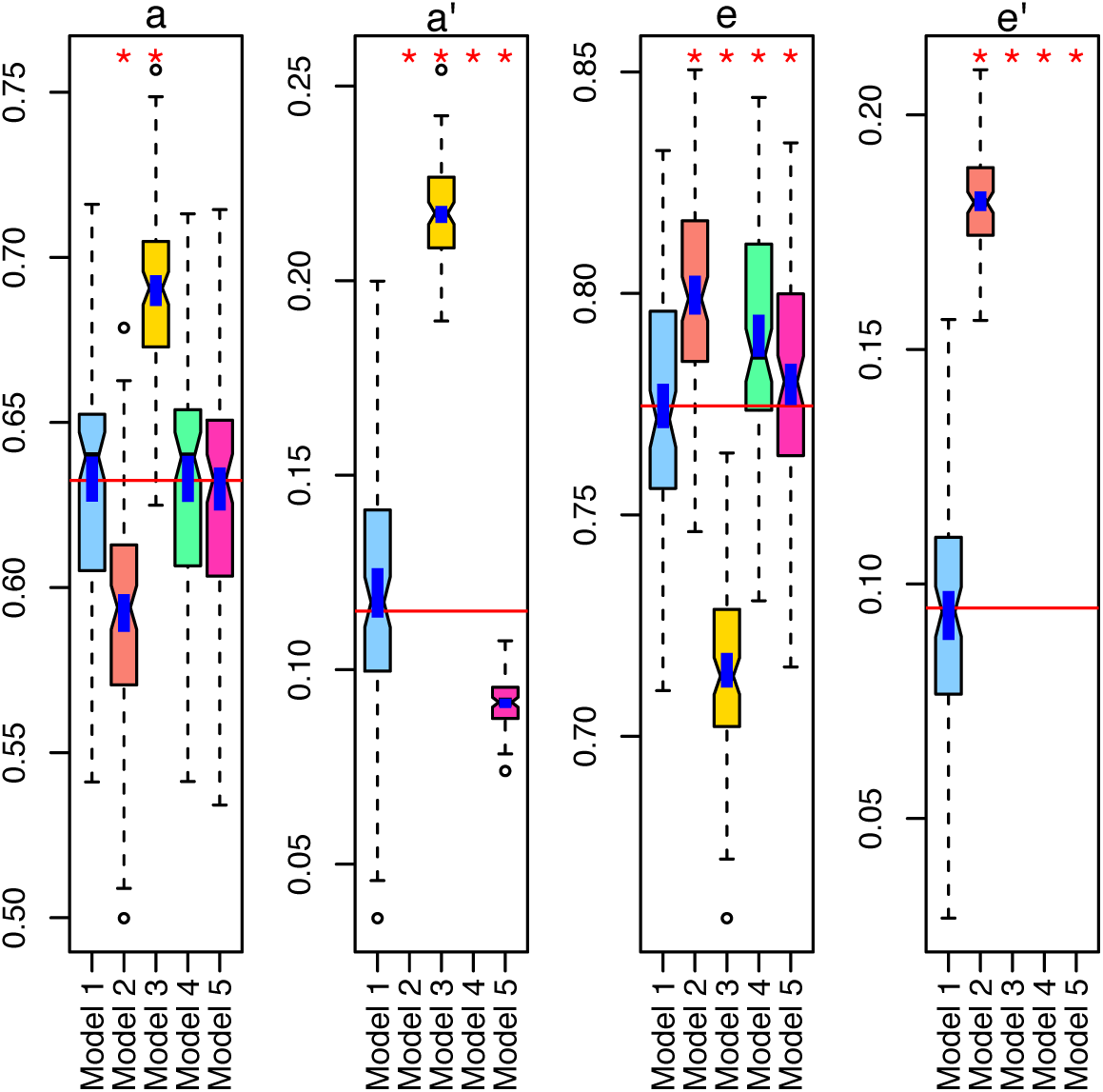
Data were simulated from model 1, and estimated with 5 different model. The vertical red lines are the true values. A red star appears above each boxplot if the estimated parameter is significantly different from its true value. The vertical blue liens are 95% confidence intervals. Model 1 is the only model that can estimate all the parameters, accurately.

It is important to note that environmental effect may be correlated with the genetic effect on the trait (*r*_*GE*_) rather than modifying the genetic effects on the trait (G×E). *r*_*GE*_ implies that certain alleles are over-or under-represented depending on the value of the moderator, and can appear as quantitative G×E in certain ways of modeling G×E, e.g., by stratifying the sample by the moderator. Entering the moderator in the means model as a main effect, as is done here, will effectively remove from the covariance model any genetic effects that are shared between trait and moderator [8].

The use of unrelated samples with genome-wide SNP data allow investigations of quantitative G×E hypotheses in larger sample sizes and on more phenotypes than are available in twin and family datasets while avoiding potential biases that exist when close relatives are modeled. To demonstrate our approach, we investigate the moderation of variance components of IQ as a function of a measure of SES (the reverse-scaled Townsend Deprivation Index) in the UK Biobank, given that this has been a hypothesis of great interest (Table S1). The estimated parameters along with their 95% confidence intervals (CI) and p-values for all five models are presented in Table 1. Across all models, the estimated parameters *a* and *e* are similar, showing consistency of the estimated V_A_ and V_E_ at the mean level of SES, and estimates of *a*′ and *e*′ are negative, showing that estimated V_A_ and V_E_ decrease as a function of SES in the range of SES investigated. Constraining the heritability to be the same across the moderator by setting *e*′ = *ea*′/*a* (Model 5 vs. Model 1) led to a non-significant decrease in fit (*p* = .145), suggesting that overall V_*p*_ changes as a function of SES and that V_A_ and V_E_ change roughly proportionately. Consistent with this, Model 5 had the lowest AIC and BIC values, making it the most parsimonious model (see Figure 3).

**Table 1:** Estimated parameters for the proposed 5 models and Turkeimer model. Green lines are CI and number in brackets are p-value. The bold numbers are the minimum AIC/BIC. Computational time and memory used in gigabyte are also reported.

| Parameter | Model |  |  |  |  |  |
| --- | --- | --- | --- | --- | --- | --- |
|  | 1 | 2 | 3 | 4 | 5 | Turkeimer |
| $\hat{\lambda}$ | -0.110 | -0.110 | -0.110 | -0.110 | -0.110 | 0.360 |
| $\hat{a}$ | 0.507 | 0.506 | 0.508 | 0.507 | 0.507 | 0.572 |
|  | (0.48,0.53) | (0.48,0.53) | (0.49,0.53) | (0.49,0.53) | (0.49,0.53) | — |
| $\hat{a}'$ | -0.026 [0.032] | — | -0.041 [1.4e-10] | — | -0.011 | 0.141 |
|  | (-0.05,0.00) | — | (-0.054,-0.028) | — | (-0.02,-0.00) | — |
| $\hat{e}$ | 0.848 | 0.849 | 0.847 | 0.849 | 0.849 | — |
|  | (0.84,0.86) | (0.84,0.86) | (0.83,0.86) | (0.84,0.86) | (0.84,0.86) | — |
| $\hat{e}'$ | -0.010 [0.108] | -0.024 [3e-10] | — | — | — | — |
|  | (-0.03,0.00) | (-0.03,-0.01) | — | — | — | — |
| Log-like | -19234.13 | -19235.85 | -19234.9 | -19254.82 | -19234.69 | -2873.2 |
| AIC | 38476.27 | 38477.71 | 38475.8 | 38513.64 | **38475.38** | 6777.3 |
| BIC | 38510.67 | 38503.51 | 38501.60 | 38530.84 | **38501.19** | 6812.5 |
| Time | 21:30:54 | 9:39:27 | 8:50:18 | 12:27:43 | 13:13:52 | — |
| Memory (GB) | 167.4 | 155.4 | 155.4 | 155.4 | 167.4 | — |

**Figure 3.**
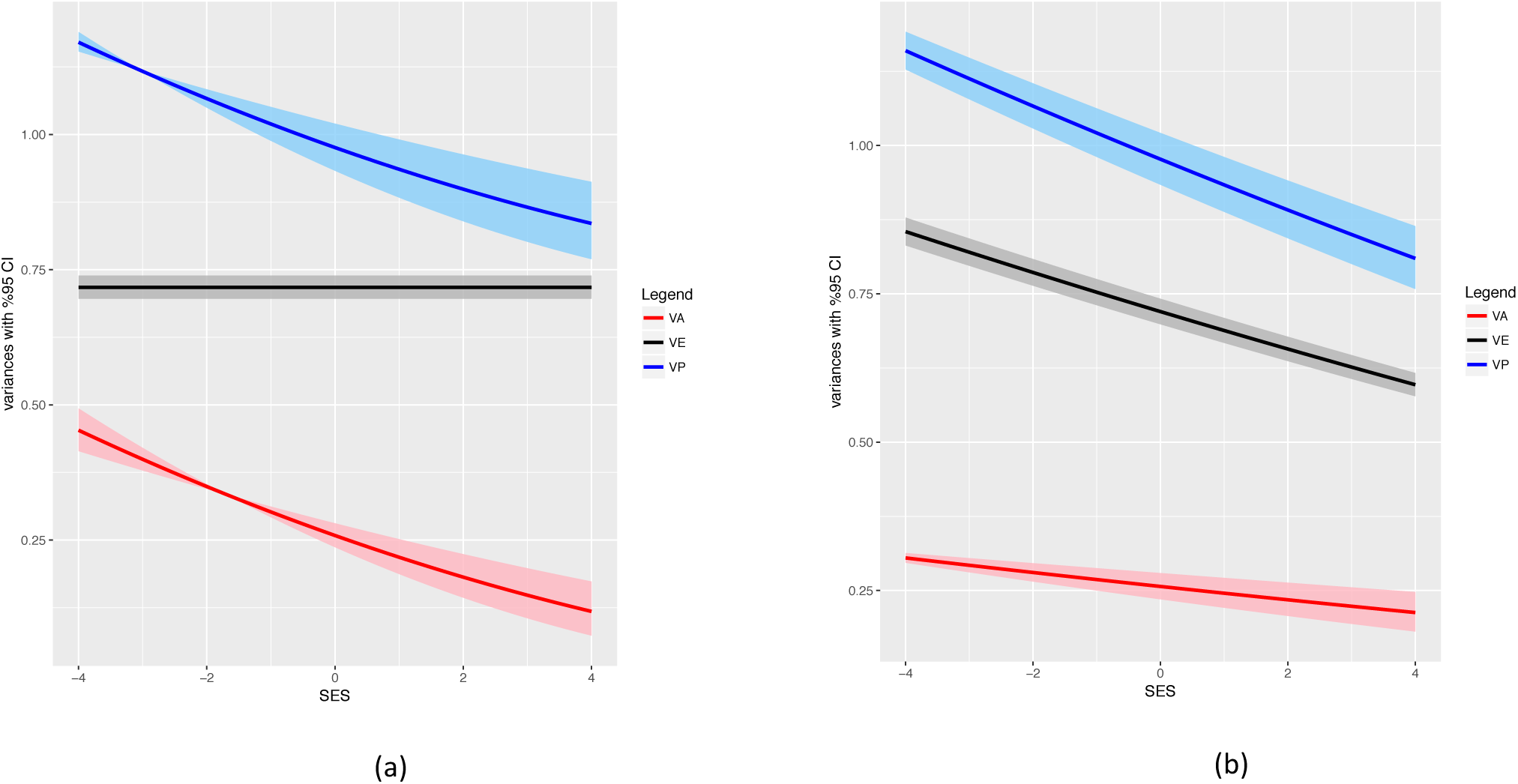
a) V_A_ = (*a* + *a*′ SES)^2^, V_E_ = *e*^2^ and V_P_ = V_A_ + V_E_ as a function of SES with a %95 CI for model 3. b) V_A_ = (*a* + *a*′ SES)^2^, V_E_ = (*e* + *ea*′/*a* SES)^2^ and V_P_ = V_A_ + V_E_ as a function of SES with a %95 CI for model 5.

Our results are in the opposite direction of those reported by several studies conducted in the US [19] [21] [22]. However, our results are more consistent with several findings from Western Europe and Australia, where V_A_, on average, decreases slightly as a function of SES [20]. However, even studies from Western Europe and Australia have tended to find virtually no change in overall V_*p*_ (due to a counter-balancing effect of V_E_ increases as a function of SES), whereas we found a significant decrease in V_*p*_ across SES. While it is possible that moderation of unmodeled non-additive genetic effects in twin studies could lead to discrepancies between the current results and those based on twins, this cannot explain different patterns of changes in V_*p*_. Thus, the source of discrepancies across this studies and previous ones based on twins may have to do with differences in measures of IQ, of SES, or in differences in study populations. Almost all the US twin studies are conducted in adolescent and early childhood, while this study and [20] are on adults (see Table S1).

There are two limitations regarding the modeling approach for quantitative G×E we introduced. First, because codes were written in R, the computational speed is not optimal (see Table 1), although we have partially resolved this problem by finding a better starting point from moment matching methods (Haseman-Elston regression). Second, we have not yet developed methods to estimate quantitative G×E for categorical outcomes, such as occurs in case-control studies. Both issues are potentially addressable with further refinement of the code and model in the future.

We have demonstrated a general approach for estimating quantitative G×E in unrelated samples using constrained optimization. We showed by simulation that the bias of the estimated parameters is negligible, that type-I errors are appropriately controlled, and that estimates can be biased under model misspecification. In particular, if quantitative G×E effects occur, we showed that traditional approaches that do not model G×E underestimate heritability. We applied our method to whole-genome SNP data from the UK Biobank, and found that phenotypic variance of IQ decreases as a function of SES, but that heritability of SES remains roughly constant.

## Materials and Methods

### Data

UK Biobank recruited 500,000 people aged between 40-79 years in 2006-2010 from across the UK. Prospective participants were invited to visit an assessment center, at which they completed an automated questionnaire and were interviewed about lifestyle, medical history and nutritional habits; basic variables such weight, height, blood pressure etc. were measured; and blood and urine samples were taken, and DNA was extracted from blood. Genotyping was done using two closely related arrays, with each having ~800,000 SNP markers. Samples were analyzed in batches of approximately 4700 individuals.

### Data quality control

Participants were tested for fluid intelligence at up to three separate occasions; when more than one score was available for an individual, we selected the first score. Fluid intelligence score is a simple unweighted sum of the number of correct answers given to the 13 fluid intelligence questions. Participants who did not answer all of the questions within the allotted 2-minute limit were scored as zero for each unanswered question. The mean for standardized fluid intelligence score was .046 in males and -.042 in females (*p* < 0.001). Participant age (mean=58.2, SD=7.99) was computed from the appropriate fluid intelligence collection date minus the birthday. The standardized fluid intelligence score decreased slightly as age increased (beta = -.0095, *p* ~ 0, adjusted R^2^ = 0.006). Townsend deprivation index (TDI) was calculated immediately prior to participant joining UK Biobank based on the area in which their postcode was located. The mean for standardized TDI was 0.0028 and –0.0025 in males and females, respectively (*p*= 0.60). In this paper, we used reverse-coded TDI as a measure for SES.

After merging fluid intelligence scores with non-missing TDI, sex, age at recruitment, place born and genotype measurement batch, 41,908 Caucasian individuals remained with genotype information. After dropping individuals with SNP missingness > .03 and dropping a minimal number of individuals in pairs with genomic relatedness > 0.05, the final sample size was 40,172. We used the first 15 principal components as covariates (see Table S25). In addition to the UKB standard genotypic quality control, we dropped SNPs with missingness > .05 and with Hardy-Weinberg equilibrium threshold *p* < 10^‒6^, leaving 345,767 SNPs.

### Simulation procedure

We simulated populations with sizes *N* ∈ {500,1000,2000,4000,8000} for different sets of parameters *θ* = (*a*, *a*′, *e*, *e*′). These values for *θ* are shown in Table S2. For *θ*_6_ with (*a*, *a*′) = (0.633,0.038), the genotypic variance was *V*_*A*_= *a*^2^ = .4 for *M* = 0 and was (*a* + *a*′*M*)^2^ = .45 for *M* = 1, i.e., increasing one standard unit of the moderator led to an increase of 0.05 units of . Similarly, for (*e*, *e*′) = (0.774, 0.032), the non-genotypic variance was *V*_*E*_ = *e*^2^ = .6 for *M* = 0 and was (*e* + *e*′*M*)^2^ = .65 for *M* = 1.

Genotypes were simulated from UK Biobank array data and phenotypes were simulated using models 1–5 for different sets of 1000 causal variants (CVs) in each replication. For each set of the parameters, we simulated *r* = 200 replications (for N=8000, *r* = 100). To gauge the performance of the proposed method, we estimated the parameters for each replication and computed the bias and variance of each estimate as

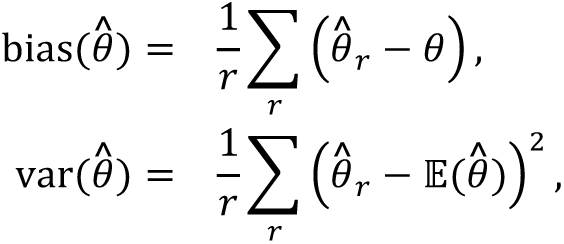

for *θ* ∈ {*a*, *a*′ , *e*, *e*′}, where **E**(*θ̂*) = 1/*r* Σ_*r*_ *θ̂*_*r*_ (Tables 2 and S3-S6).

**Table 2.** Simulation results for model 1 with different set of parameters and sample sizes. Columns are bias and standard deviation of the estimated parameters.

| True Parameters |  |  |  | N | a |  | a' |  | e |  | e' |  |
| --- | --- | --- | --- | --- | --- | --- | --- | --- | --- | --- | --- | --- |
| a | a' | e | e' |  | bias | sd | bias | sd | bias | sd | bias | sd |
| 0.633 | 0.115 | 0.775 | 0.095 | 2000 | -0.009 | 0.113 | 0.003 | 0.073 | -0.009 | 0.088 | 0.003 | 0.061 |
|  |  |  |  | 4000 | -0.005 | 0.059 | -0.002 | 0.051 | -0.001 | 0.045 | 0.004 | 0.040 |
|  |  |  |  | 8000 | 0.000 | 0.035 | 0.004 | 0.033 | 0.000 | 0.025 | -0.002 | 0.027 |
| 0.633 | 0.077 | 0.775 | 0.095 | 2000 | -0.015 | 0.110 | -0.001 | 0.075 | -0.011 | 0.090 | -0.004 | 0.058 |
|  |  |  |  | 4000 | -0.003 | 0.058 | -0.001 | 0.057 | -0.005 | 0.046 | 0.003 | 0.046 |
|  |  |  |  | 8000 | -0.001 | 0.030 | -0.001 | 0.034 | -0.003 | 0.022 | 0.001 | 0.027 |
| 0.633 | 0.077 | 0.775 | 0.063 | 2000 | -0.004 | 0.105 | 0.000 | 0.073 | -0.014 | 0.084 | 0.002 | 0.059 |
|  |  |  |  | 4000 | -0.003 | 0.052 | 0.003 | 0.060 | -0.002 | 0.040 | 0.001 | 0.047 |
|  |  |  |  | 8000 | 0.000 | 0.033 | 0.008 | 0.034 | -0.002 | 0.024 | -0.006 | 0.027 |
| 0.633 | 0.077 | 0.775 | 0.032 | 2000 | -0.001 | 0.115 | -0.006 | 0.080 | -0.019 | 0.090 | 0.004 | 0.066 |
|  |  |  |  | 4000 | 0.001 | 0.054 | -0.010 | 0.056 | -0.007 | 0.042 | 0.007 | 0.047 |
|  |  |  |  | 8000 | 0.003 | 0.031 | -0.002 | 0.037 | -0.004 | 0.022 | 0.002 | 0.029 |
| 0.633 | 0.038 | 0.775 | 0.063 | 2000 | -0.012 | 0.125 | -0.004 | 0.075 | -0.011 | 0.089 | 0.004 | 0.058 |
|  |  |  |  | 4000 | -0.003 | 0.062 | 0.004 | 0.061 | -0.006 | 0.048 | -0.003 | 0.049 |
|  |  |  |  | 8000 | -0.001 | 0.035 | 0.003 | 0.038 | -0.003 | 0.026 | -0.002 | 0.030 |
| 0.633 | 0.038 | 0.775 | 0.032 | 2000 | -0.022 | 0.117 | -0.004 | 0.077 | 0.000 | 0.081 | 0.004 | 0.062 |
|  |  |  |  | 4000 | -0.006 | 0.058 | -0.005 | 0.065 | -0.002 | 0.047 | 0.005 | 0.053 |
|  |  |  |  | 8000 | 0.000 | 0.034 | 0.004 | 0.035 | -0.002 | 0.024 | -0.003 | 0.027 |

To investigate statistical power and type-I error rates, we simulated r data sets with sizes *N* ∈ {500,1000,2000,4000,8000} from models defined under the alternative and null hypotheses respectively, and then computed the maximum value of the log-likelihood for the alternative, *ℓ*(*Θ*_H_) and the null, *ℓ*(*Θ*_0_). The test statistics is

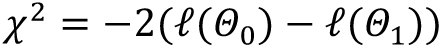

was compared to the critical value *Q*_1‒*α*_, which is obtained from the central chi-square distribution with *df* degrees of freedom and *α* = 0.05, where *df* is the difference between the number of free parameters of models alternative and null. Power was computed as

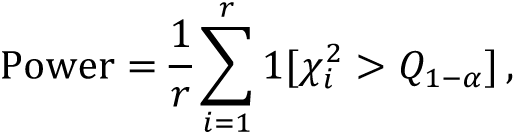

where *r* = 200 replications. Similarly, we computed the type I error by simulating *r* data sets from the null distribution, and calculated the proportion of rejected test,

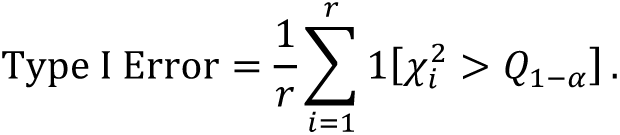

### Code availability

The R codes are freely available at https://github.com/rtahmasbi/G×E.

## Acknowledgements

This work was supported by the National Institutes of Mental Health grant R01MH100141 to Dr. Keller. We thank Jian Yang and Peter Visscher for helpful comments on the project.

## Author contributions

R.T. and M.K. designed the study. E.T. discussed the results and commented on the final manuscript. All authors wrote the manuscript.

## Additional information

**Supplementary information** is available for this paper.

